# Chronotherapy as a Potential Strategy to Reduce Ifosfamide-Induced Encephalopathy: A Preclinical Study in a Murine Model

**DOI:** 10.1101/2025.08.28.672634

**Authors:** Mylene Malek Chennoufi, Dorra Dridi, Khaled Lasram, Nouha Ben Abdeljalil, Asma Omezzine, Benoit Mauvieux, Yvan Touitou, Ahmed Sleheddine Masmoudi, Naceur A. Boughattas

## Abstract

**Purpose:** Nearly five decades after its introduction into clinical oncology, ifosfamide (IFO) remains a widely used alkylating chemotherapeutic agent for the treatment of sarcomas, germ-cell tumors and selected hematological malignancies in both adult and pediatric oncology. Its use is limited by severe toxicities, particularly hemorrhagic cystitis and central nervous system complications. Recent studies have highlighted the persistence and complexity of IFO-related neurotoxicity (**Idle & Beyoğlu, 2023 [1])**, including an increased risk of encephalopathy associated with certain formulations in pediatric populations (**Hillaire-Buys et al., 2019 [2]**) and differences between originator and generic formulations (**Chambord et al., 2023 [3]**), these concerns prompted a regulatory safety review by the **E**uropean **M**edicines **A**gency **[4]**. Circadian rhythms are increasingly recognized as key regulators of drug metabolism and toxicity. In a previous chronotolerance study, we demonstrated a marked time-of-day–dependent variation in lethal IFO toxicity at the LD50 dose (**Chennoufi & Boughattas, 2025 [5]**). This study aimed to determine whether sublethal IFO exposure (LD30) also exhibits circadian variation in multi-organ toxicity.

**Methods:** One hundred male Swiss albino mice synchronized under a 12:12 h light–dark cycle received IFO LD30 at four circadian times (1, 7, 13 and 19 hours after light onset, HALO). Hematological parameters, hepatic enzymes, histopathology and neurobehavioral performance were evaluated.

**Results:** Toxicity varied significantly according to dosing time. Administration at 7 HALO produced the most pronounced injury, whereas dosing at 13 HALO was associated with reduced hepatic, renal and bladder damage. Encephalopathy-related lesions were predominantly observed at 19 HALO.

**Conclusion:** Sublethal IFO toxicity exhibits clear circadian variation. Administration at 13 HALO corresponded to the previously identified peak of LD50 tolerance, supporting further exploration of time-adjusted ifosfamide administration strategies to improve treatment tolerability.

## Introduction

Nearly five decades after its introduction into clinical oncology, ifosfamide (IFO) remains a widely used alkylating chemotherapeutic agent for the treatment of sarcomas, germ-cell tumors and selected hematological malignancies in both adult and pediatric oncology (**Zalupski & Baker, 1988 [6]**; **Williams & Wainer, 1999 [7]**; **Furlanut & Franceschi, 2003 [8]**; **Dechant et al., 1991[9]**), however, its clinical use continues to be limited by severe toxicities, particularly hemorrhagic cystitis and central nervous system complications such as ifosfamide-induced encephalopathy. Despite improvements in supportive care, these adverse effects remain difficult to predict and prevent, resulting in a persistently narrow therapeutic index. Recent clinical and pharmacological studies have highlighted the persistence and complexity of IFO-related neurotoxicity (**Idle & Beyoğlu, 2023 [1]; Beyoğlu D, 2025 [10]**), including increased encephalopathy risk associated with certain formulations in pediatric populations **(Hillaire-Buys et al., 2019 [2])** and differences between originator and generic formulations **(Chambord et al., 2023 [3])**. These concerns have led to a regulatory safety evaluation by the European Medicines Agency in 2023 **[4]**, emphasizing the need for improved strategies to enhance treatment tolerability.

In parallel, circadian rhythms are increasingly recognized as key regulators of drug metabolism, detoxification pathways and tissue susceptibility. Consequently, the timing of drug administration has emerged as a critical determinant of both efficacy and toxicity in oncology. While chronotherapy has been explored for several anticancer agents, the circadian determinants of ifosfamide toxicity remain insufficiently characterized, particularly at the level of organ-specific injury (**Lévi 1997 [11]; Lévi & Schibler, 2010 [12]; Fu & Kettner, 2013 [13])**

In oncology practice, several trials have re-examined IFO-based regimens, including combinations with immunotherapy, regional hyperthermia, and anthracycline-based protocols, showing that IFO remains a cornerstone of current sarcoma management rather than a drug of the past **(El-Tanani M, 2024 [14])**. Together, these observations confirm that IFO is undergoing a real renaissance, both clinically and scientifically.

Although chronotherapy of cyclophosphamide has been documented **(Haus et al., 1974 [15])**, the circadian determinants of IFO toxicity remain poorly understood, despite its complex metabolism via multiple cytochrome P450 isoenzymes and the recognised vulnerability of the brain to its toxic metabolites.

In a previous chronotolerance study, we demonstrated a marked time-of-day–dependent variation in lethal IFO toxicity at the LD50 dose, identifying a clear temporal pattern of susceptibility and tolerance (**Chennoufi & Boughattas, 2025 [5]**). However, the use of lethal dosing prevented detailed investigation of organ-specific toxic effects. The present study was therefore designed to determine whether sublethal ifosfamide exposure (LD30) also displays circadian variation in multi-organ toxicity. Using a controlled experimental model, we evaluated hematological, hepatic, urothelial and neurobehavioral endpoints across four circadian dosing times (HALO), with the aim of identifying the temporal window associated with reduced global toxicity and providing a translational basis for optimized ifosfamide chronotherapy.

## Methods

### Animals and Synchronization

A total of 100 male Swiss albino mice (SIPHAT, Tunis), aged 8–10 weeks (≈25 g body weight), were used in the present study. Animals were synchronized under a controlled 12:12 h light–dark cycle in accordance with established chronobiological standards for experimental design (**Touitou et al., 2004 [16]; Refinetti R2004 [17])**. Animals were housed at room temperature 22 ± 2 °C, food and water ad libitum). Two separate rooms were used to avoid night-time manipulation: Room 1 (lights on at 7:00 a.m.) and Room 2 (lights on at 8:00 p.m.). Experimental time points are expressed as Hours After Light Onset (HALO), with HALO 0 defined as the time of light onset in each room. Circadian entrainment was verified by monitoring daily oscillations in core body temperature prior to experimentation.

### LD50 and LD30 Determination

LD50 and LD30 values were previously determined by probit analysis in our chronotolerance study (Chennoufi MM & Boughattas NA, 2025). The estimated LD50 was approximately 520 mg/kg and the sublethal LD30 approximately 450 mg/kg.

### LD30 Acute Toxicity Protocol

To assess organ-specific toxicity while minimizing lethality, mice received a single intraperitoneal injection of IFO at the LD30-equivalent dose (≈450 mg/kg). Injections were administered at four predefined time points: 1, 7, 13, and 19 HALO. All animals were scheduled for sacrifice 48 hours after injection at the same HALO as dosing to maintain temporal alignment. No longitudinal survival follow-up was performed, as survival was not a predefined endpoint of the present study. Any mortality occurring prior to the planned endpoint was recorded. Animals were randomly allocated to treatment groups. Sample size per group is specified in each figure legend.

### Endpoints Measured

Blood samples were collected for hematological parameters (WBC, lymphocytes, RBC, platelets) and biochemical markers (ALT, AST). Organs (liver, kidney, bladder, brain) were harvested for histopathological examination with blinded scoring. Neurological function was assessed using the wire suspension test immediately prior to sacrifice.

### Hematotoxicity Evaluation

Animals were sacrificed 48 hours after IFO administration by decapitation in accordance with approved ethical procedures. Blood was collected immediately into appropriate collection tubes for hematological and biochemical analyses. Complete Blood Counts (CBC) were performed using an automated hematology analyzer.

Parameters assessed:

- White blood cell count (WBC, 10^3^/mm^3^)
- Platelet count (PLT, 10^3^/mm^3^)
- Red blood cell count (RBC, 10□/mm^3^)
- Lymphocytes (10^3^/mm^3^)

### Hepatotoxicity Evaluation

Serum was obtained by centrifugation and analyzed for hepatic transaminases:

- Aspartate transaminase (AST, IU/L)
- Alanine transaminase (ALT, IU/L)

Enzyme activity was determined using commercial colorimetric kits (Spinreact, Spain) according to the manufacturer’s instructions.

### Histopathology

Organs were immediately fixed in 10% buffered formalin, embedded in paraffin, sectioned at 5 µm, and stained with hematoxylin–eosin (H&E). Lesions were scored semi-quantitatively on a scale from 0 to 5 (0 = no lesion; 5 = severe damage) by a pathologist blinded to treatment group and dosing time. Organ-specific criteria included:

- Bladder: mucosal erosion, hemorrhage, edema
- Liver: hepatocyte necrosis, inflammation, fatty change
- Brain: neuronal degeneration, edema, vacuolization
- Kidney: tubular epithelial degeneration, cytoplasmic vacuolization, interstitial edema, inflammatory infiltration

### Neurotoxicity Functional Test (Wire Test)

Neurobehavioral performance was assessed 48 hours post-treatment using the wire suspension test. Each mouse was placed on a horizontal wire (2 mm diameter, 40 cm length, 30 cm above a padded surface). Latency to fall (maximum 60 seconds) was recorded.

### Statistical Analysis

Data are presented as mean ± SEM. Continuous variables were analyzed using two-way ANOVA with Treatment (Control vs LD30) and Dosing Time (1, 7, 13, 19 HALO) as independent factors, including the Treatment × Time interaction term. When appropriate, post hoc comparisons were performed using Tukey’s test. Histopathological scores were analyzed using the Kruskal–Wallis test followed by Dunn’s post hoc test. Statistical significance was set at p < 0.05. Analyses were performed using GraphPad Prism (version X, GraphPad Software, USA).

## Results

### 1. Animal Synchronization

### 2. Hematotoxicity

Data are presented as mean ± SEM. **Two-way ANOVA** demonstrated a significant effect of dosing time **treatment, and their interaction (Treatment × Time) on all hematological parameters**, indicating a strong circadian modulation of IFO-induced hematotoxicity (Fig. 2 A, B, C, D).

**Fig. 1:**
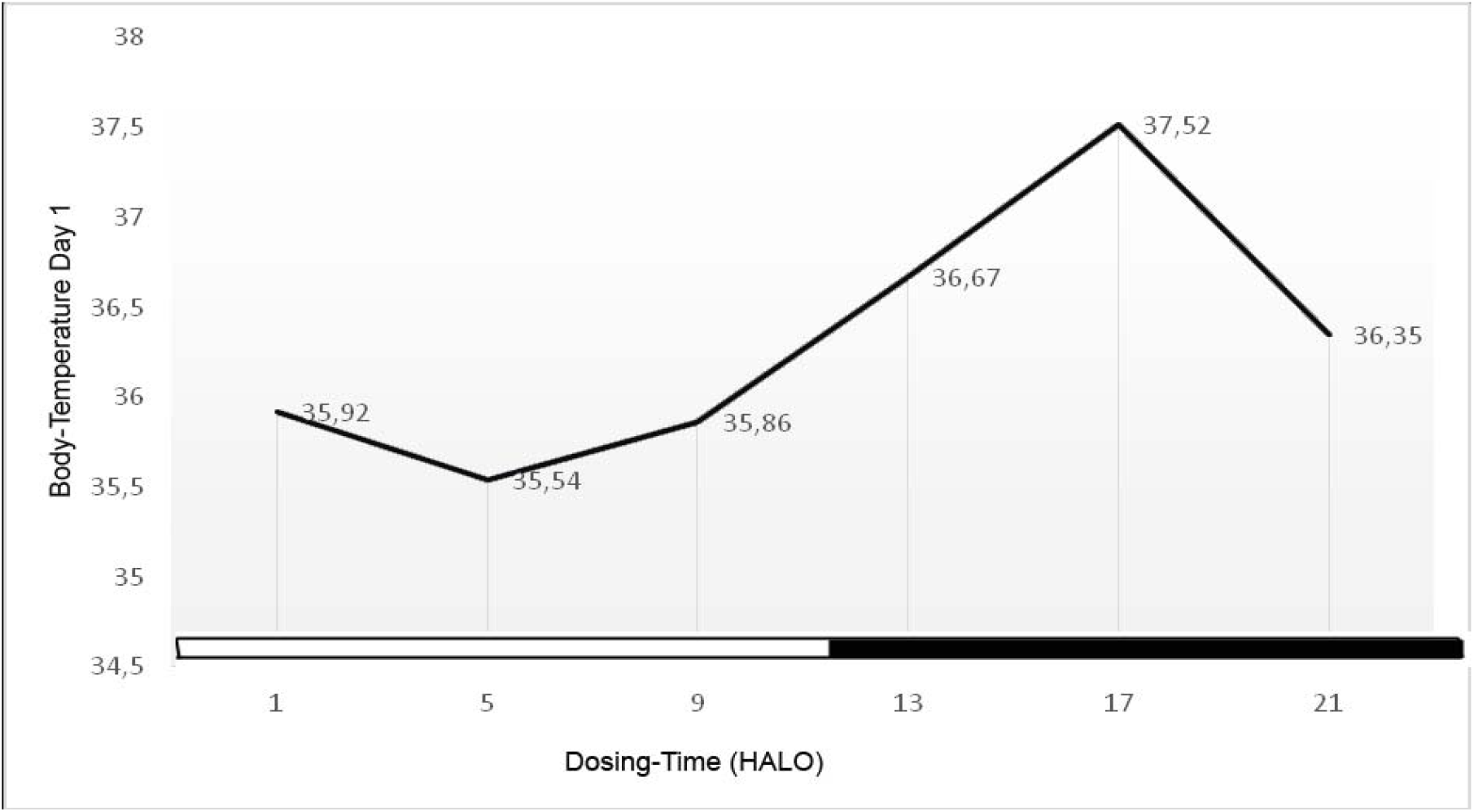
Circadian variation of rectal temperature in mice before Ifosfamide administration. Animals were maintained under a 12:12 h light–dark cycle. Daily oscillations in core body temperature consistent with circadian entrainment were observed prior to treatment, in agreement with both our previous chronotolerance study (Fig. 1) and established reference patterns **(Waterhouse et al., 2005 [18])**.

**Fig. 2.**
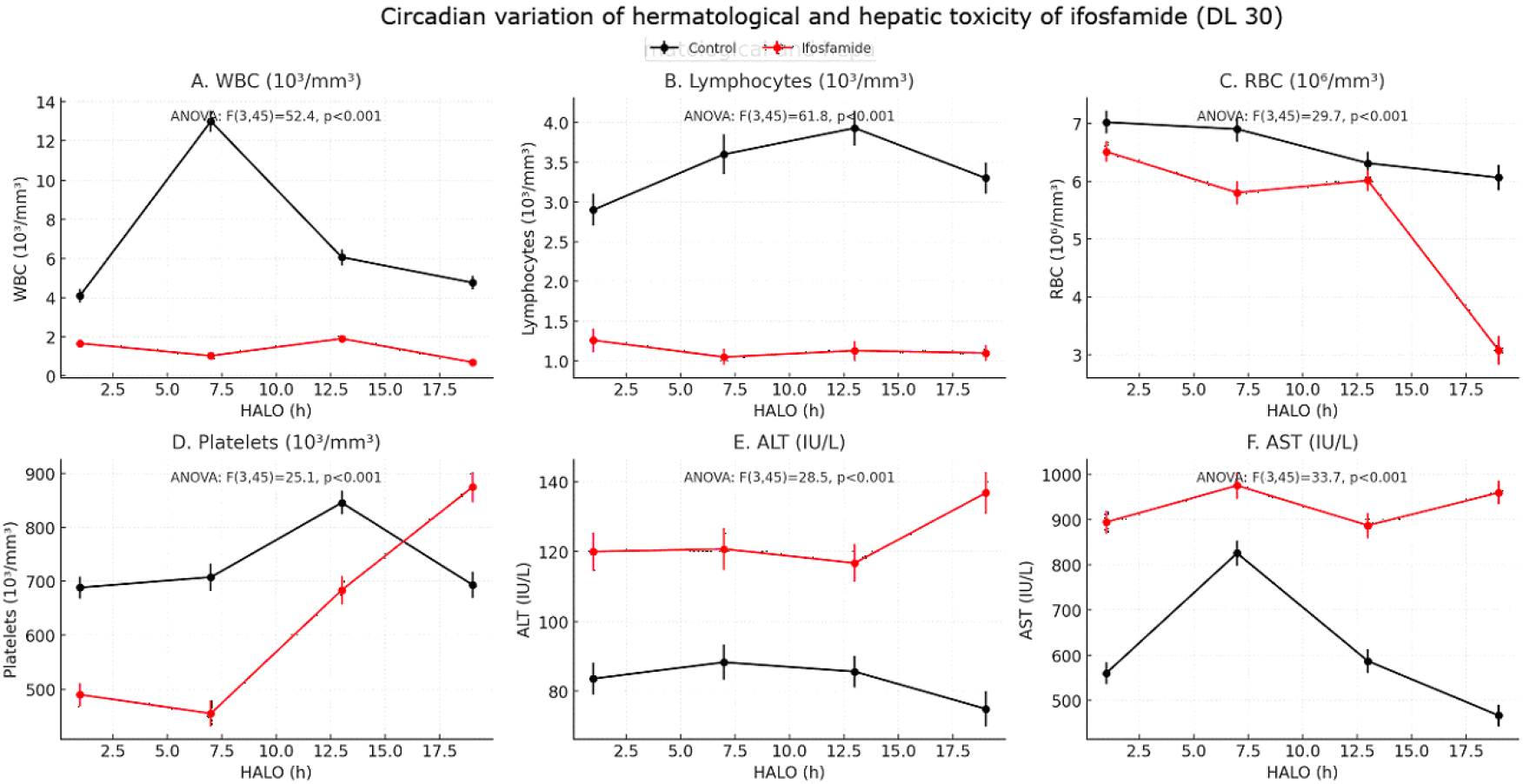
Circadian variation of hematological and hepatic toxicities of ifosfamide (LD30). **(A)** White blood cells (WBC, 10^3^/mm^3^): Ifosfamide induced a marked leukopenia at all dosing times, most pronounced at 7 and 19 HALO. **(B)** Lymphocytes (10^3^/mm^3^): Severe lymphopenia was observed, with significant circadian variation and nadirs at 7 and 19 HALO. **(C)** Red blood cells (RBC, 10^3^/mm^3^): A profound decrease was recorded at 19 HALO compared with other times. **(D)** Platelets (10^3^/mm^3^): Thrombocytopenia was significant at 7 HALO, partial recovery occurred at 13 HALO, while paradoxical thrombocytosis appeared at 19 HALO. **(E)** Alanine aminotransferase (ALT, IU/L) and **(F)** Aspartate aminotransferase (AST, IU/L)

A significant circadian variation was observed for all endpoints (WBC: F(3,45) = 52.4; lymphocytes: F(3,45) = 61.8; RBC: F(3,45) = 29.7; platelets: F(3,45) = 25.1; all p < 0.001) (Fig. 2A–D).

### White Blood Cells (WBC)

IFO induced marked leukopenia at all dosing times. The most pronounced suppression occurred at 7 and 19 HALO, whereas partial preservation was observed at 13 HALO compared to time-matched controls (Fig. 2A).

### Lymphocytes

Lymphocyte counts showed a similar pattern, with the greatest suppression at 7 and 19 HALO and relatively higher values at 13 HALO (Fig. 2B).

### Red Blood Cells (RBC)

RBC levels were significantly reduced at 19 HALO, while values at 1 and 13 HALO were closer to control levels (Fig. 2C).

### Platelets (PLT)

Thrombocytopenia was most evident at 7 HALO. Platelet counts at 13 HALO were comparatively preserved, whereas 19 HALO was associated with paradoxical thrombocytosis (Fig. 2D).

Overall, hematotoxicity was more pronounced when IFO was administered at 7 and 19 HALO, whereas 13 HALO was associated with comparatively lower hematological alterations.

## 3. Hepatic Toxicity

Transaminase levels peaked at 7 and 19 HALO, whereas comparatively lower elevations were observed at 13 HALO relative to time-matched controls (Fig. 2E–F).

These findings indicate a clear time-of-day dependence of early hepatic injury following LD30 administration. Hepatic enzyme elevations displayed significant circadian dependency, peaking at 7 and 19 HALO, whereas 13 HALO showed relative protection. ALT: F(3,45) = 28.5, p < 0.001; AST: F(3,45) = 33.7, p < 0.001.

## 4. Histopathology

Semi-quantitative lesion scores revealed a significant effect of dosing time across all organs (Kruskal–Wallis test), indicating circadian variation in tissue susceptibility.

Bladder: χ^2^(3) = 14.8, p = 0.002

Liver: χ^2^(3) = 18.2, p < 0.001

Brain: χ^2^(3) = 9.7, p = 0.045

Kidney: χ^2^(3) = 11.6, p = 0.021

Histopathological assessment confirmed organ-specific differences according to dosing time (Fig. 3). Bladder and liver were the primary targets of toxicity. Lesion severity scores varied significantly across dosing times (Kruskal–Wallis test).

**fig. 3.**
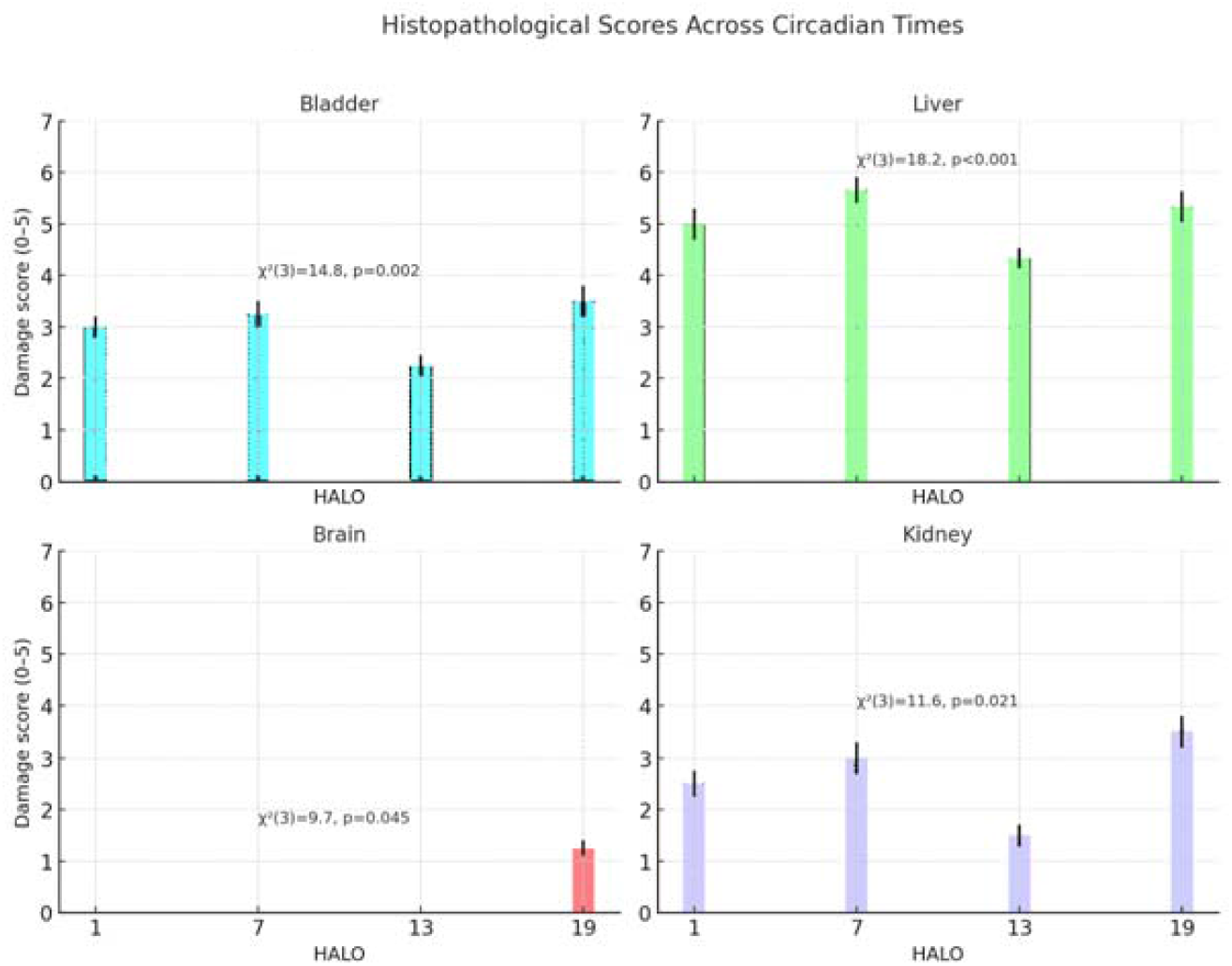
Circadian variation of ifosfamide-induced histopathological lesions in mice. Semi-quantitative damage scores (mean ± SEM, for bladder, liver, kidney, and brain across four dosing times (1, 7, 13, and 19 HALO). (A) bladder; (B) liver; (C) brain (D) kidney

The highest bladder damage was observed at 19 HALO.

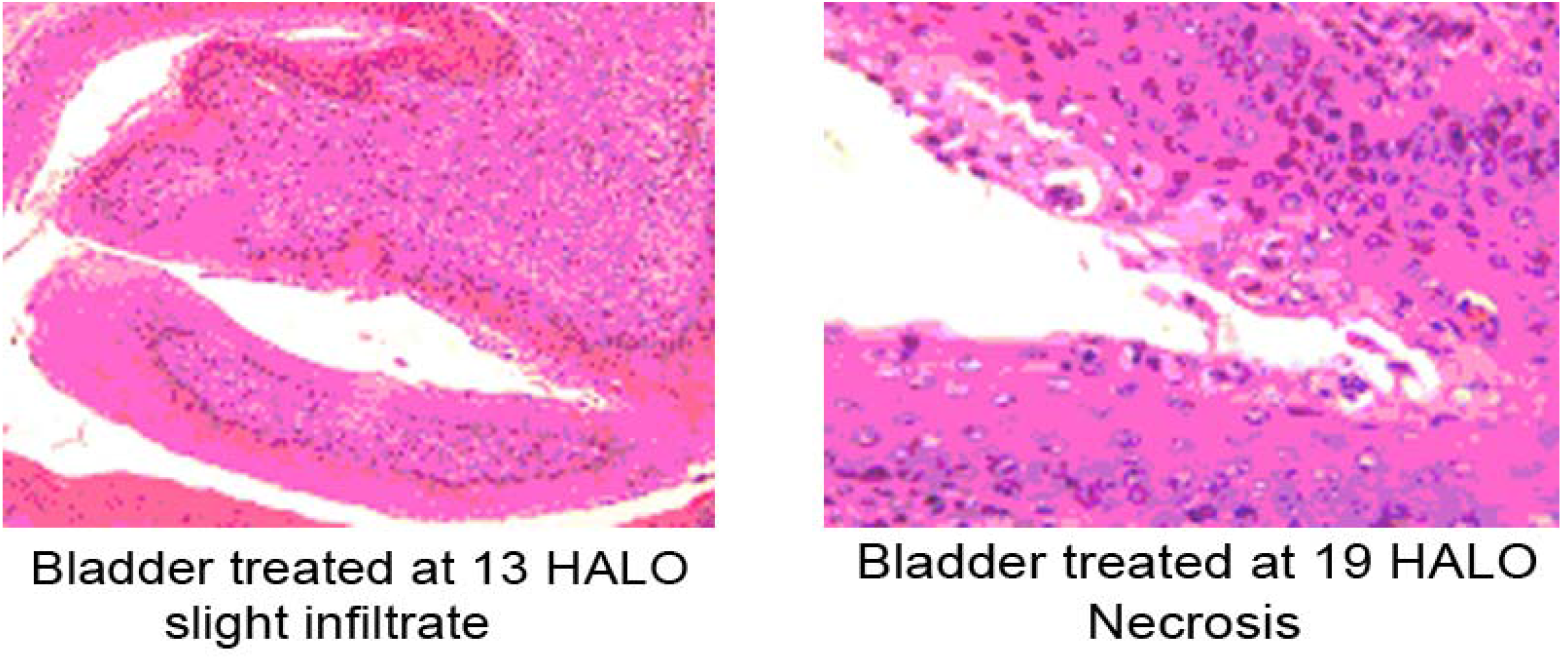

Hepatic necrosis and inflammatory infiltration were most pronounced at 7 HALO and comparatively less severe at 13 HALO.

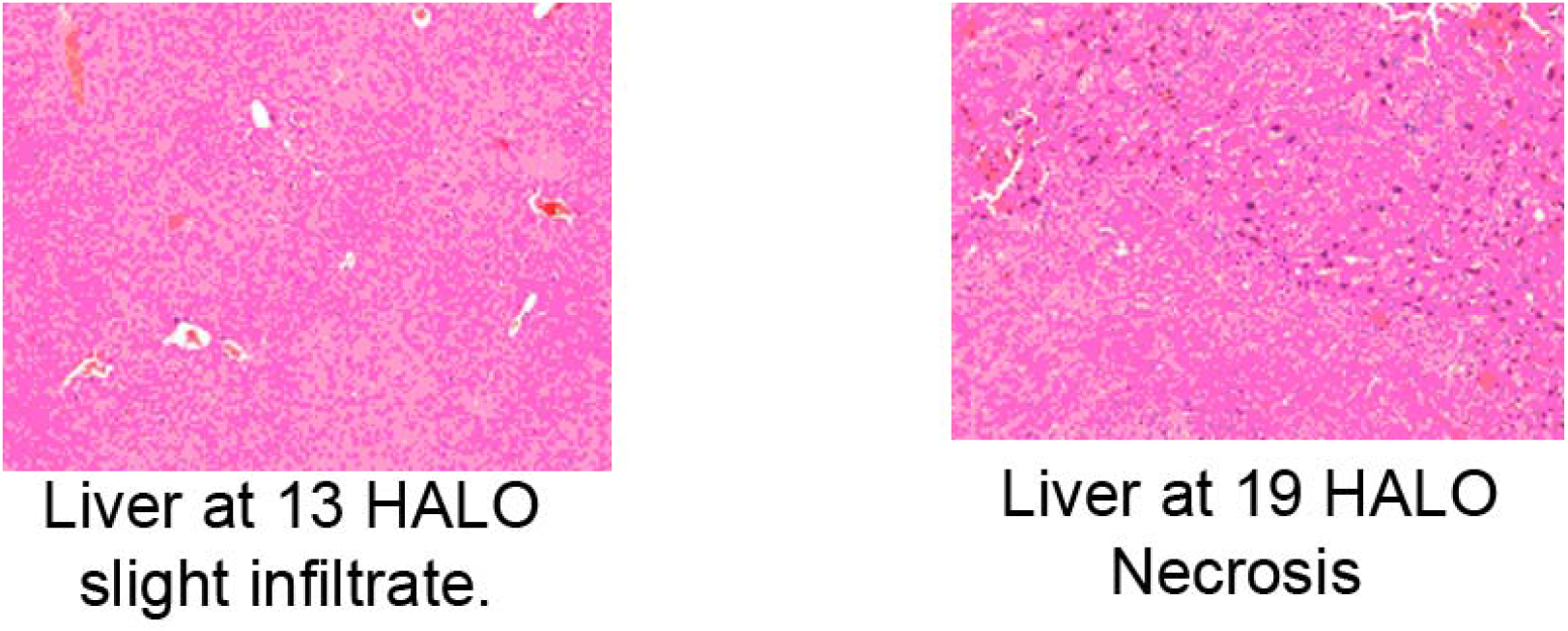

Brain lesions were minimal at most time points but increased at 19 HALO.

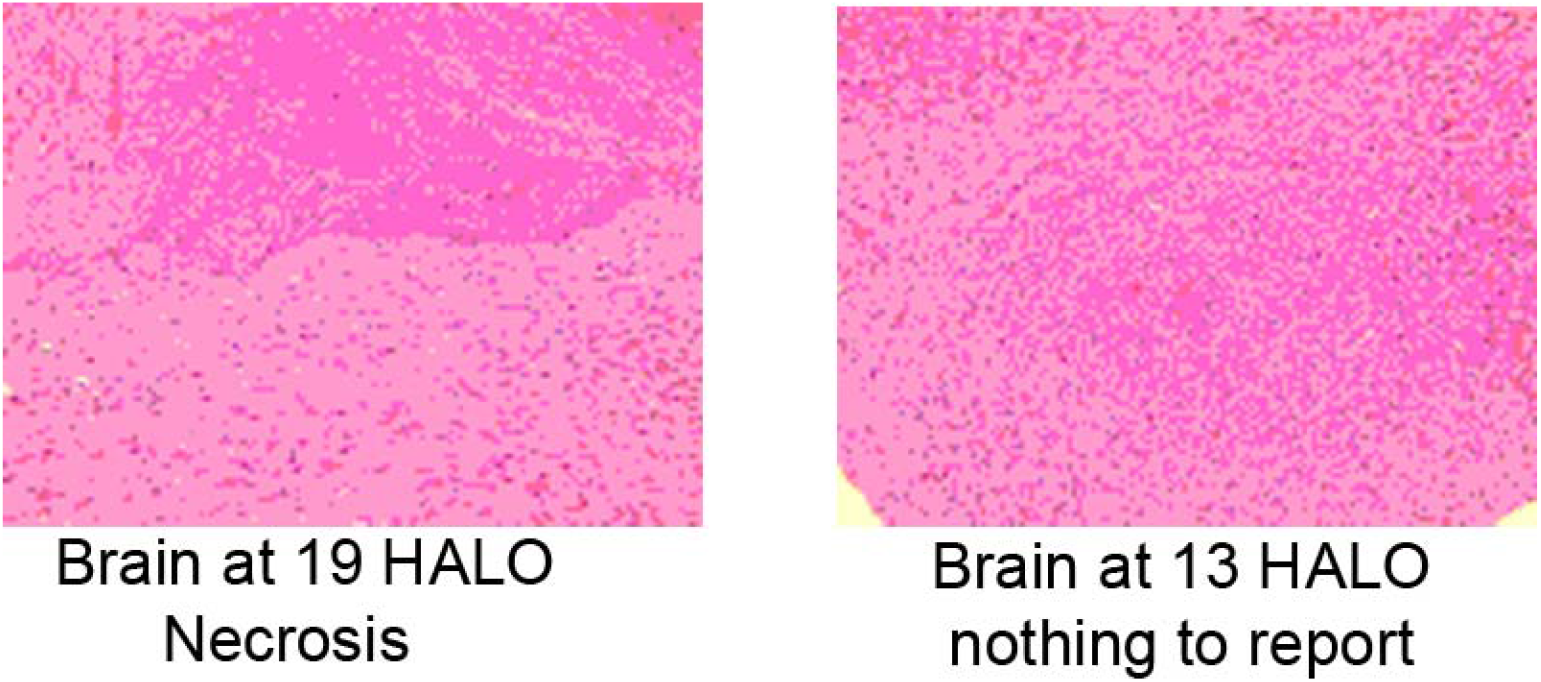

Kidney injury also demonstrated dosing-time dependence, with highest scores at 19 HALO and near-baseline values at 1 and 13 HALO.

These results indicate differential organ vulnerability depending on the timing of IFO administration.

## 5. Neurotoxicity (Wire Test)

Neurobehavioral performance differed according to dosing time (two-way ANOVA, significant Treatment × Time interaction). The shortest latency to fall, indicating maximal impairment, was observed at 19 HALO, whereas performance at 13 HALO was comparatively preserved relative to treated groups at other time points (Fig. 4).

**Fig. 4.**
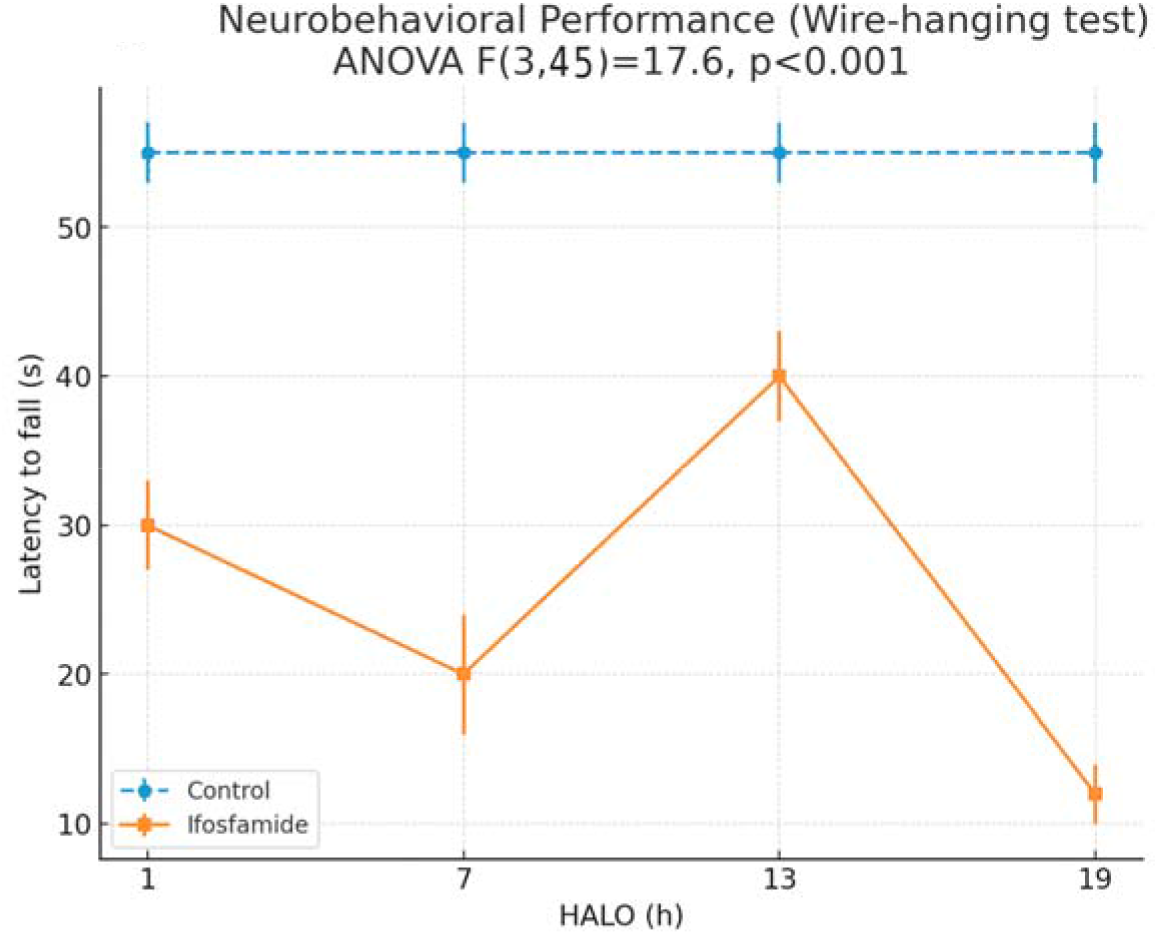
Neurobehavioral performance in the wire-hanging test after ifosfamide administration (LD30) in mice. Latency to fall (mean ± SEM, n = 6–8 per group) was recorded at 1, 7, 13, and 19 HALO. Ifosfamide-treated mice exhibited significant circadian variation in neuromuscular coordination (ANOVA F (3,45) = 17.6, p < 0.001). The shortest latency to fall, indicating maximal impairment, was observed at 19 HALO (∼12 s), while the longest latency was at 13 HALO (∼40 s), suggesting partial preservation of motor performance. Control mice maintained stable latencies (∼55 s) across all time points.

## Discussion

The present study demonstrates that Ifosfamide (IFO) toxicity exhibits a marked circadian variation across multiple organs—including the liver, kidney, bladder and brain—as well as hematological and neurobehavioral endpoints. These findings extend our previous LD50 chronotolerance study, which established a robust circadian rhythm in IFO lethality, with optimal tolerance at 13 HALO and maximal vulnerability around 7 and 19 HALO. Here, using a sublethal LD30-equivalent dose, we show that acute multi-organ toxicity follows the same temporal pattern.

Hematological alterations revealed a strong time-of-day dependence. Leukopenia and lymphopenia were most pronounced at 7 and 19 HALO, whereas counts were better preserved at 1 and 13 HALO. Platelets also exhibited a circadian pattern, with significant thrombocytopenia at 7 HALO, partial recovery at 13 HALO, and paradoxical thrombocytosis at 19 HALO. These variations may reflect the circadian rhythmicity of bone marrow proliferation, immune cell trafficking, and the temporal regulation of drug metabolism pathways. Hepatic toxicity showed a similar rhythmic pattern. Both ALT and AST peaked at 7 HALO, consistent with histopathological findings of marked hepatocellular necrosis and inflammation. The lowest enzyme elevations and histological damage were observed at 13 HALO, suggesting that hepatic detoxification capacity is maximally aligned with IFO metabolism during this phase.

Renal and urothelial injuries also varied significantly across the circadian cycle. Bladder damage, including erosions, hemorrhage and edema, was most severe at 19 HALO, whereas kidney lesions (tubular degeneration, vacuolization and interstitial edema) peaked at the same time point. These converging results are consistent with the susceptibility of renal tubular cells to chloroacetaldehyde, a toxic IFO metabolite known to follow circadian variation in formation.

Brain toxicity was relatively mild except at 19 HALO, where neuronal degeneration and vacuolization became prominent. This increased vulnerability coincided with the poorest performance in the wire-hanging test, indicating a functional correlate of histopathological damage. The alignment of metabolic stress, toxic metabolite accumulation, and reduced antioxidant capacity during this phase may contribute to heightened neurological sensitivity.

These results are particularly relevant in view of recent publications questioning the consistency of Ifosfamide-induced encephalopathy. In their review entitled *“Ifosfamide-induced encephalopathy: Myth or Reality?”*, **Abahssain H** et al. **[19]**, highlighted the heterogeneity of clinical presentations and the absence of reliable predictive markers. Our findings provide experimental evidence that neurological vulnerability to IFO is **not a myth**, but a **real, measurable, and time-dependent phenomenon**. The marked increase in neuronal lesions and motor impairment at 19 HALO strongly supports the existence of a circadian window of susceptibility, offering a physiological explanation for the variability observed in patients.

Taken together, these findings demonstrate that IFO-induced toxicity is not constant throughout the 24-h cycle but instead follows a reproducible circadian pattern. The optimal tolerance window consistently occurred at **13 HALO**, reinforcing the notion that aligning IFO administration with circadian physiology could significantly reduce adverse effects. This concept is consistent with earlier chronotherapy studies on other alkylating agents, including cyclophosphamide.

The translational implication is that timing optimization may improve the therapeutic index of IFO in clinical settings, particularly given the drug’s narrow safety margin and variability in neurotoxicity. Future clinical studies are warranted to determine whether administering IFO at patient-specific circadian phases can reduce encephalopathy, urotoxicity and systemic toxicity while maintaining antitumor efficacy.

### Clinical implications

Taken together, these results highlight that IFO toxicity is not only dose-dependent but also time-dependent. The **maximum toxicity at 7 and 19 HALO** underscores the need to avoid dosing during these circadian phases. Conversely, **13 HALO emerged as the most protective time**, consistent with our earlier survival study. For humans, this corresponds approximately to the beginning of the **daytime activity period**, reinforcing the rationale for daytime administration in clinical protocols (Table 1).

**Table 1.**
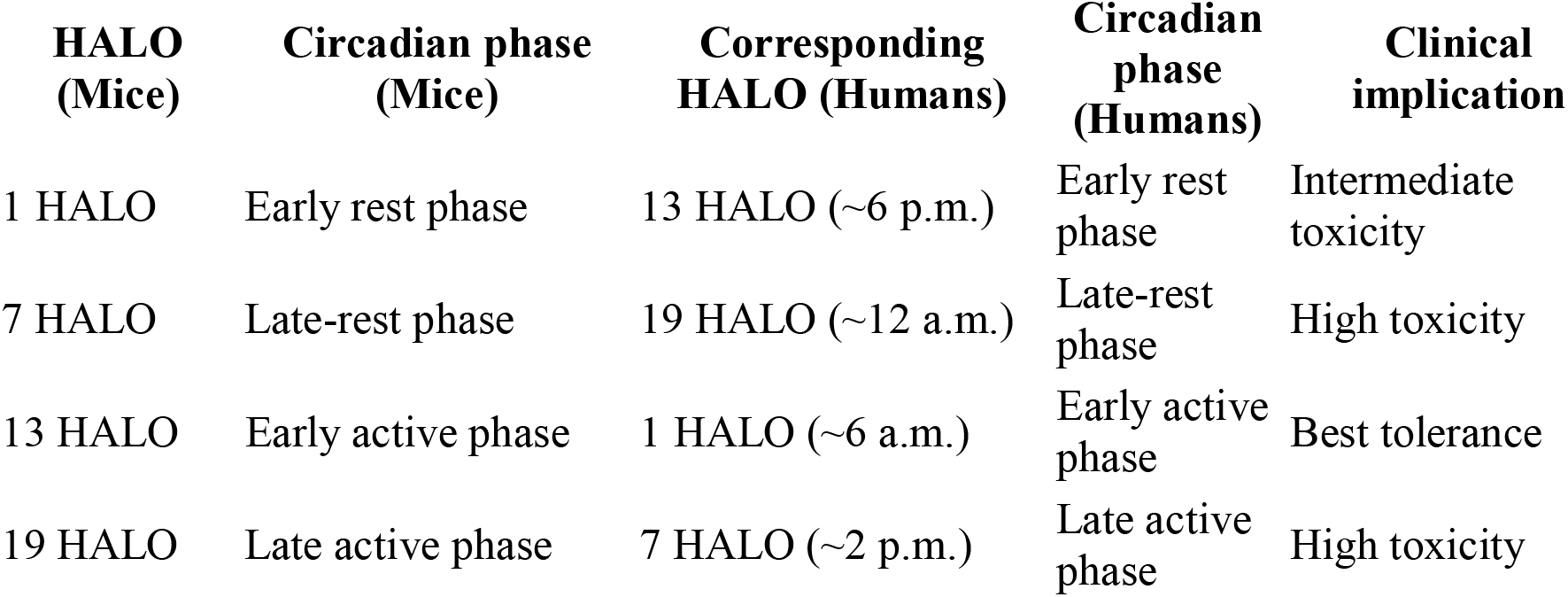
Correspondence between circadian dosing time (HALO) in mice and humans, after LD30 ifosfamide in mice.

While mesna reduces urotoxicity and methylene blue can mitigate encephalopathy in some cases, no strategy fully prevents IFO’s major toxicities. Our findings suggest that **chronotherapy may provide an additional, non-pharmacological approach to improve tolerability**, allowing safer administration of IFO in oncology, particularly in pediatric protocols where it remains widely used.

## Conclusion

This study demonstrates that ifosfamide (IFO) exhibits marked **circadian variation in organ-specific toxicity**. Hematological suppression, hepatic enzyme elevation, urothelial lesions and neurotoxicity were all significantly influenced by dosing time. The most severe toxic effects occurred at **7 and 19 HALO**, while the **least toxicity was consistently observed at 13 HALO**. These findings extend our previous work on IFO chronotolerance by highlighting the **specific targets of circadian toxicity**—bone marrow, liver, bladder and brain pathways. Importantly, neurological impairment, a hallmark of ifosfamide-induced encephalopathy (IIE), was minimized when treatment was given at 13 HALO. Given that IIE remains unpredictable and lacks an effective preventive or curative strategy, chronotherapy emerges as a promising, non-pharmacological approach to improve IFO tolerability. Optimizing administration schedules could reduce systemic and organ-specific toxicities, thereby enhancing the therapeutic index of IFO in clinical oncology, particularly in pediatric protocols where it continues to play a crucial role.

## Declarations

### Author contributions

**MMC** designed the study, performed the experiments, analyzed the data, and wrote the manuscript. **DD** contributed through scientific advice. **KL** assisted with animal dissections. **NBA** performed the anatomical pathology analyses. **AO** carried out the hematological analyses. **BM** contributed to data analysis. **YT** critically reviewed the manuscript. **ASM** reviewed the present manuscript. **NAB** supervised the study. All authors approved the final version.

### Data availability

The datasets generated and analyzed during this study are available from the corresponding author on reasonable request.

### Ethics

All procedures involving animals were conducted in accordance with the international guidelines for the care and use of laboratory animals. The experimental protocol was reviewed and approved by the Institutional Animal Ethics Committee of the Faculty of Medicine of Monastir.

## Acknowledgements

This study was carried out at the Laboratory of Pharmacology, Faculty of Medicine of Monastir, under the supervision of Prof. Naceur A. Boughattas. The authors sincerely thank him for his scientific guidance throughout the project. We thank as well the Fattouma Bourgiba University Hospital and Sahloul University Hospital Staff for support and dedicated assistance in animal handling and laboratory facilities.

## References

1. Idle JR, Beyoğlu D. Ifosfamide – History, efficacy, toxicity and encephalopathy. Pharmacol Ther. 2023. doi:10.1016/j.pharmthera.2023.108366

2. Hillaire-Buys D, Mousset M, Allouchery M, et al. Liquid formulation of ifosfamide increased risk of encephalopathy: A case-control study in a pediatric population. Therapies 2019. 10.1016/j.therap.2019.08.001

3. Chambord J, Henny F, Salleron J, et al. IfosfamideLJinduced encephalopathy: BrandLJname (HOLOXAN®) vs generic formulation (IFOSFAMIDE EG®). J Clin Pharm Ther. 2019 10.1111/jcpt.12823

4. European Medicines Agency (EMA). Ifosfamide solutions – referral: Benefits of ifosfamide solutions continue to outweigh risks. 2021. Reference Number: EMA/219444/2021 https://www.ema.europa.eu/en/medicines/human/referrals/ifosfamide-solutions

5. Chennoufi MM, Boughattas NA. Chronotolerance of ifosfamide in mice: evidence for a circadian rhythm. Chronobiol Int. 2025. doi: 10.1080/07420528.2025.2581095.

6. Zalupski M, Baker LH. Ifosfamide. J Natl Cancer Inst. 1988. doi:10.1093/jnci/80.8.556.

7. Williams M, Wainer IW. Cyclophosphamide versus ifosfamide: to use ifosfamide or not to use, that is the three-dimensional question. Curr Pharm Des. 1999. doi:10.2174/1381612993399953.

8. Furlanut M, Franceschi L. Pharmacology of Ifosfamide. Oncology. 2003. doi:10.1159/000073350

9. Dechant KL, et al. Ifosfamide/mesna: a review of its antineoplastic activity, pharmacokinetic properties and therapeutic efficacy in cancer. Drugs. 1991. doi:10.2165/00003495-199142030-00006

10. Beyoğlu D, Hamberg P, IJzerman NS, Mathijssen RHJ, Idle JR. New metabolic insights into the mechanism of ifosfamide encephalopathy. Biomed Pharmacother. 2025. doi: 10.1016/j.biopha.2024.117773.

11. Lévi F. Circadian chronotherapy for human cancers. Lancet Oncol. 1997

12. Lévi F, Schibler U. Chronotherapeutics: the relevance of timing in cancer therapy. Cancer Causes Control. 2010;21(2):201–210. doi:10.1007/s10552-009-9476-8.

13. Loning Fu, Kettner NM. The circadian clock in cancer development and therapy. Prog Mol Biol Transl Sci. 2013; doi:10.1016/B978-0-12-396971-2.00009-9.

14. El-Tanani M, Rabbani SA, Ali AA, Alfaouri IGA, Al Nsairat H, Al-Ani IH, Aljabali AA, Rizzo M, Patoulias D, Khan MA, Parvez S, El-Tanani Y. Circadian rhythms and cancer: implications for timing in therapy. Discov Oncol. 2024. doi: 10.1007/s12672-024-01643-4.

15. Haus E, et al. Murine circadian susceptibility rhythm to cyclophosphamide. Chronobiologia. 1974.

16. Touitou Y, et al. Ethical principles and standards for the conduct of human and animal biological rhythm research. Chronobiol Int. 2004. doi:10.1081/CBI-120030045.

17. Refinetti R. Variability of diurnality in laboratory rodents. Chronobiol Int. 2004;21(1):191–198.

18. Waterhouse J, et al. The circadian rhythm of core temperature and physiological implications. Chronobiol Int. 2005;22(2):207–225.

19. Abahssain H, Moukafih B, Essangri H, Mrabti H, Meddah B, Guessous F, Fadhil FZ, Souadka A, Errihani H. Methylene blue and Ifosfamide-induced encephalopathy: Myth or Reality?. J Oncol Pharm Pract. 2021 Jan;27. doi: 10.1177/1078155220971843.

